# CURCUMIN HAS LOWER IC50 VALUES AGAINST A549 LUNG CANCER CELLS

**DOI:** 10.1101/2023.08.06.552182

**Authors:** Sedat Kaçar, Özlem Tomsuk, Varol Şahintürk

## Abstract

In this study, we aimed to determine the toxic effect of curcumin on A549 lung cancer cells and to show how this effect is reflected in morphology. Firstly, the toxic doses of curcumin against A549 lung cancer cells were determined via MTT and neutral red cytotoxicity tests by using the doses of 6.25, 12.5, 25, 50, 100, 200 and 400 µM. The morphological examination was performed using inverted microscope and light microscope with hematoxylin-eosin eosin.As a result, the IC50 dose of curcumin against A549 cells was 33 µM according to MTT cytotoxicity test and 52 µM according to the neutral red cytotoxicity test. When these doses were administered, the rounded and shrunken cells were visible in the inverted microscope, while apoptotic hallmarks such as nucleus condensation, renal nucleus structure and cellular shrinkage were detected in hematoxylin-eosin staining. In conclusion, the low IC50 values of curcumin for 24 hours indicate that curcumin is effective against A549 cells. It has the potential to be used either alone or in combination with other agents in cancer studies in case its low toxic effect on normal was assured.

## 1. Introduction

Curcumin, extracted from Turmeric rhizomes (*Curcuma longa)*, or diferuloylmethane with its chemical name is a yellow polyphenol having broad range of beneficial activities, including, anti-inflammatory, anti-oxidant and anti-carcinogenic activities [1, 2]. Anti-carcinogenic effects of curcumin were demonstrated several cancer cells and tissues [3-6].

As a result of studies of curcumin’s effect on cancer cells, relatively lower half inhibitory concentrations (IC50) have been obtained, suggesting potency of curcumin to be used in cancer treatment alone or with other less toxic agents. In literature as a drawback of curcumin, its less solubility in aqueous media is highlighted. To circumvent this issue, studies such as water-soluble nanoparticle to find out different remedies are ongoing [7]

Of most rampant cancers, lung cancer is the forefront, with its high mortality rate in the world. A549 cells are a commonly utilized model cell line for lung cancer studies as well as for toxicity studies regarding lung metabolism and carcinogenesis [8]. In this respect, herein, we plan to investigate anti-proliferative and cancer cell morphology deforming properties of curcumin on A549 cells by 2 cytotoxicity test and microscopic examination.

## 2. Materials and Methods

### 2.1 MTT assay

MTT, used for determining cell proliferation, measures the mitochondrial enzyme activity of live cells and is based on the conversion of tetrazolium dye (purple formazan) to determine cell viability [9]. A549 human lung cancer cells were seeded on to 96-well plates at a density of 5×10^3^ cells per well. After the incubation period of 24 hours, the cells were treated with decreasing concentrations of curcumin from 400 µM to 0 (6.3, 12.5, 25, 50, 100, 200, 400 µM) for 24 hours. Subsequently, MTT solution diluted in pre-prepared medium was added to each well and incubated at least 2 hours. After that, 100 µl of DMSO was added to each well to dissolve the formazan crystals. The cell viability was quantified by ELISA reader at 570 nm. The results were calculated relative to the viability of control (vehicle-treated) cells, of which the viability percentage was accepted as 100% and which were treated with only 0.1 %. DMSO.

### 2.2 Neutral red test

Neutral red uptake assay is one of the common on and relatively sensitive cytotoxicity tests and relied on the ability of the viable cells to incorporate the neutral red into their lysosomes In the assay, as the cells start to be non-viable, then they are able to incorporate no more neutral red. Thus, the more neutral red uptake indicates higher cell survival [10, 11]. The cytotoxicity of curcumin on A549 cells was further checked by neutral red uptake assay. A549 human lung cancer cells were seeded on to 96-well plates at a density of 5×10^3^ cells per well. After the incubation period of 24 hours, the cells were treated with decreasing concentrations of curcumin from 400 µM to 0 (6.3, 12.5, 25, 50, 100, 200, 400 µM) for 24 hours. Subsequently, 1% neutral red solution diluted in pre-prepared medium was added to each well and incubated at least 3 hours. After that, 100 μl of DESORB (50% ethanol + 1% glacial acetic acid) solution was added to each well to dissolve the neutral red crystals. The cell viability was quantified by ELISA reader at 540 nm. The results were calculated relative to the viability of control (vehicle-treated) cells, of which the viability percentage was accepted as 100% and which were treated with only 0.1 %. DMSO.

### 2.3 Inverted microscopy

The curcumin-induced morphological changes were observed under an inverted microscope. Concisely, A549 cells were grown in 6-well plates till they adhered to the flask. Then, the cell medium was disposed and washed by PBS. Later, the cells were treated with curcumin for 24 hours. The next day, the cells were observed under the microscope, and the respective images were taken.

### 2.4 Hematoxylin-eosin staining

The curcumin-induced morphological changes were scrutinized by hematoxylin-eosin stain, which is a rampant method preferred for detailed analysis of cytoplasmic, nuclear, and extracellular structures [12]. Concisely, A549 cells were grown in 6-well plates till they adhered to the flask. Then, the cell medium was disposed of the flask and washed by PBS. Later, the cells were treated with curcumin for 24 hours. The next day, the cells were fixed with ice-cold methanol for 10 min and washed with PBS to get rid of remaining methanol. Once stained with hematoxylin for 10 minutes, the cells were washed in 1% ammonia water to render them bluish color. Thereafter, the cells were stained with eosin stain for 10 minutes. Ultimately, the cells were washed with distilled water, dried and mounted with the water-based mounting solution.

## 3. Results

### 3.1 MTT results

As depicted in **Figure 2** and **Table 1**, the cell viability of A549 cells decreased to 89.3% at the lowest applied dose, 6.3 µM of curcumin (p>0.05 vs control). At the doses of 12.5 and 25.0 µM, the viability was 62.0 and 51.2%, respectively (p<0.01 and p<0.001 vs control, respectively). At the doses of 50.0 and 100.0, the viability was below 40% (p<0.001 vs control for all). Finally, at the highest applied two doses, almost no cell viability was detected (p<0.001 vs control for all).

**Figure 1:**
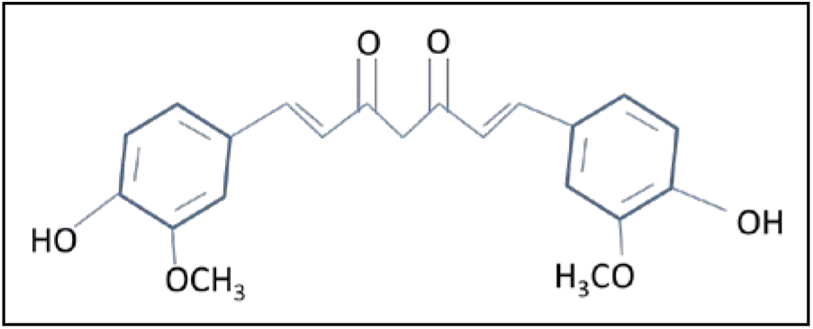
Chemical structure of curcumin

**Figure 2.**
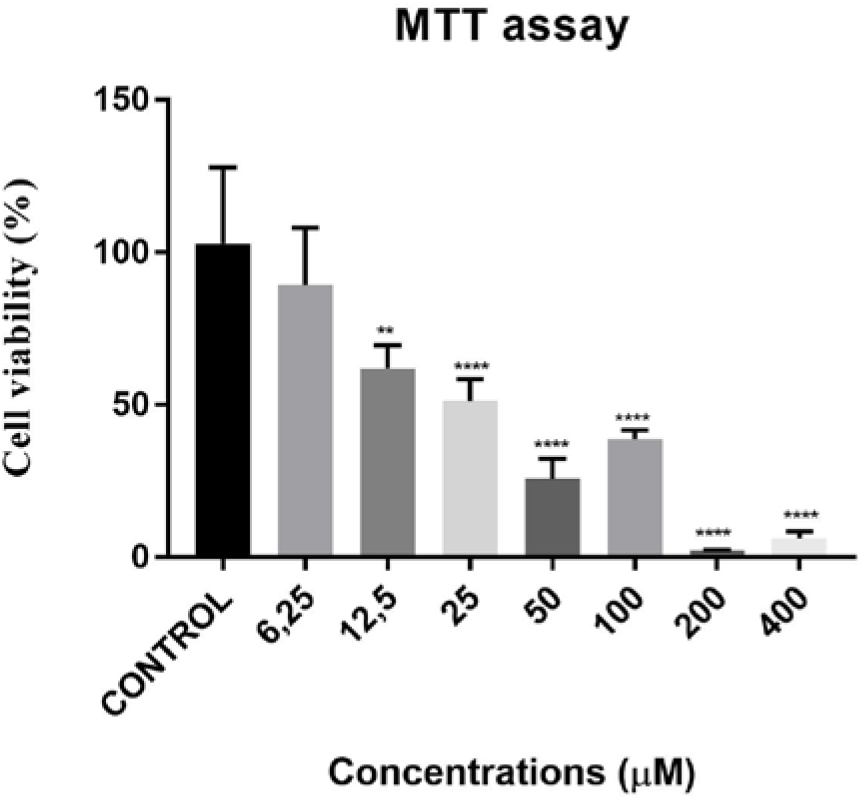
Cell viability percentages vs ascending curcumin doses according to MTT assay. *, **, *** and **** designate a respective significant difference of p>0.05, p>0.01, p>0.001 and p<0.0001 when compared to untreated cells.

**Table 1.** Numerical values of MTT and neutral red test results as a percentage.

| Curcumin doses (μM) | MTT<br>Cell viability (%)<br>(mean±SD) | NR<br>Cell viability (%)<br>(mean±SD) |
| --- | --- | --- |
| Untreated (control) | 102.9 ± 25.0 | 100.0 ± 9.9 |
| 6.3 | 89.3 ± 18.8 | 97.3 ± 13.7 |
| 12.5 | 62.0 ± 7.6 | 64.9 ± 5.6 |
| 25.0 | 51.2 ± 7.2 | 59.0 ± 10.9 |
| 50.0 | 25.7 ± 6.6 | 51.9 ± 13.4 |
| 100.0 | 38.8 ± 2.9 | 29.8 ± 12.7 |
| 200.0 | 2.1 ± 0.3 | 0.7 ± 4.1 |
| 400.0 | 6.2 ± 2.2 | 0.0 ± 0.3 |

### 3.2 Neutral red results

As depicted in **Figure 3** and **Table 1**, the cell viability of A549 cells was very similar to that of control (p>0.05 vs control). At the doses of 12.5 and 25.0 µM, the viability was 64.9 and 59.0%, respectively (p<0.001 vs control for both). At the doses of 50.0 and 100.0, the viabilities were 51.9 and 29.8% (p<0.001 vs control for both). Finally, at the highest applied two doses, almost no cell viability was detected (p<0.001 vs control for both).

**Figure 3.**
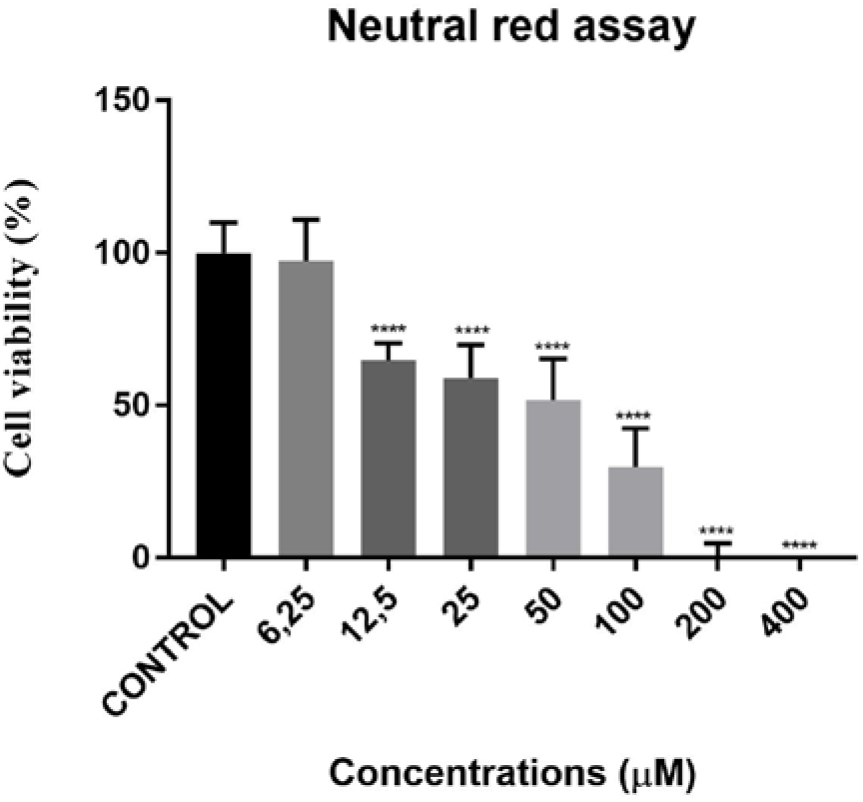
Cell viability percentages vs ascending curcumin doses according to neutral red assays. **** designates a significant difference of p<0.0001 when compared to untreated cells.

### 3.3 Inverted microscope results

In inverted microscope, the untreated lung cancer cells showed their normal morphology and were confluent all over the plate. On the other hand, curcumin-treated cells were smaller and covered less volume when compared to untreated cells. In addition, the number of cells was reduced (**Figure 4**).

**Figure 4.**
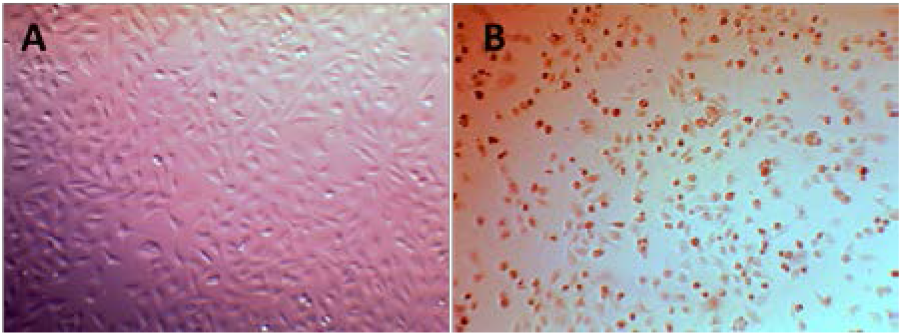
Inverted microscopy of A549 cells. **A** indicates control cells. **B** indicates curcumin-treated A549 cells. Note of the curcumin-treated cells, which are smaller and cover less volume on the plate

### 3.4 Hematoxylin-eosin results

In hematoxylin-eosin stain, the untreated lung cancer cells showed their normal morphology. However, curcumin-treated cells exhibited several abnormalities. The cells were shrunk and smaller. Besides, the typical deformation was the condensed nuclei (**Figure 5**).

**Figure 5.**
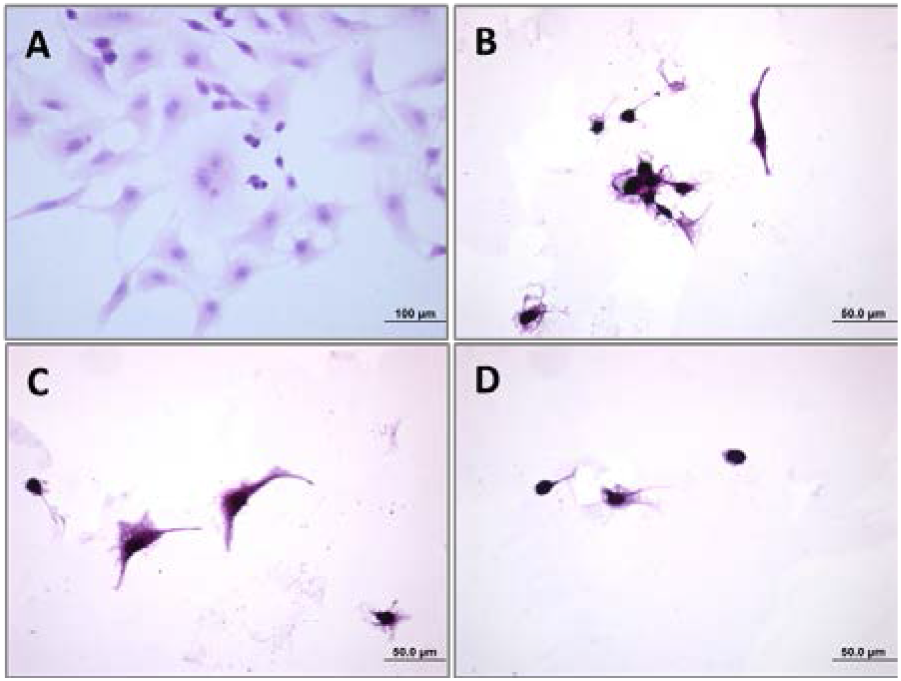
Hematoxylin-eosin staining of A549 cells. **A** indicates control cells. **B, C** and **D** indicate curcumin-treated A549 cells. Note of the abnormal and shrunk cells with condensed nuclei. Bars indicate 50 µm.

## 4. Discussion

A number of experiments were performed to reveal the anti-cancerogenic impact of curcumin in different cells or tissues [3-6], including breast cancer, nasopharyngeal carcinoma, lung cancer, colorectal cancer, etc. The goal of this research was to find out the anti-proliferative and morphological effect of curcumin on human lung cancer cells with the help of MTT and cytotoxicity assays, whereby the antiproliferative concentration of curcumin was determined, and hematoxylin-eosin staining, whereby the morphological changes were examined. Among the doses that we used here, we revealed that curcumin starts to inhibit cell proliferation effectively nearly at 12.5 µM concentration in both cytotoxicity assays. Scott and Loo (2004) used 10 and 20 µM curcumin and reported it to induce concentration-dependent DNA damage and lead to apoptotic characteristics in HCT-116 colonocytes [13]. In a recent study of Wang et al (2019), effect of curcumin on two non-small-cell lung cancer cells, A549 and SPC-A1 were studied. Curcumin doses ranging between 0-100 µM was used. In their MTT data, IC50 of curcumin against these cell lines is estimated to be between 40-50 µM [14]. Eskiler et al. (2019), in her recent article, studied breast cancer cell lines of MCF-7 and MDA-MB-231 and reported IC50 of curcumin for 48 hours as 25.6 and 8.05 µM, respectively [15].

To recap, our data substantiated the previous results in terms of the anti-proliferative effect of curcumin. Considering that the curcumin gives lower IC50 values even in the short-term studies, it can be focused to be used against the different cancers by itself or by combining with other less toxic substances.

## Acknowledgments

This study has been presented in 4nd International Material Science and Technology in Kızılcahamam (IMSTEC’19), October, 18-20, 2019, ANKARA, Turkey.

